# The effect of the menstrual cycle on the circulating microRNA pool in human plasma: a pilot study

**DOI:** 10.1101/2022.07.22.501154

**Authors:** Séverine Lamon, Joane Le Carré, Giuseppe Petito, Hong Phuoc Duong, François Luthi, Danielle Hiam, Bertrand Léger

## Abstract

**Study question:** Do ovarian hormones levels influence cf-miRNA expression across the menstrual cycle?

**Summary answer:** Measures of ovarian hormones should be rigorously included in future studies assessing cf-miRNA expression in females and used as time-varying confounders. This exploratory study suggests that cf-miRNAs may play an active role in the regulation of the female cycle in various target tissues.

**What is known already:** Cell-free or “circulating” miRNAs (cf-miRNAs) are secreted from tissues into most physiological fluids, including plasma, where they play a role in cross-tissue communication. Endogenous and exogenous factors, including sex hormones, regulate cellular miRNA expression levels. Plasma cf-miRNA levels vary with numerous pathological and physiological conditions, including in females.

**Participants/materials, setting, methods:** We conducted an exploratory study where blood samples were collected from sixteen eumenorrheic females in the early follicular phase, the ovulation phase and the mid-luteal phase of the menstrual cycle. Ovarian hormones oestrogen, progesterone, luteinizing hormone (LH) and follicle-stimulating hormone (FSH) were measured in serum by electrochemiluminescence. The expression levels of 179 plasma-enriched miRNAs were profiled using a PCR-based panel, including stringent internal and external controls to account for the potential differences in RNA extraction and reverse-transcription stemming from low-RNA input samples.

**Study design, size, duration:** This was a prospective monocentric study conducted between March and November 2021.

**Main results and the role of chance:** This exploratory study suggests that cf-miRNAs may play an active role in the regulation of the female cycle in various target tissues. Linear mixed-models adjusted for the relevant variables showed numerous associations between phases of the menstrual cycle, ovarian hormones and plasma cf-miRNA levels. Validated gene targets of the cf-miRNAs varying with the menstrual cycle were enriched within the female reproductive tissues and primarily involved in cell proliferation and apoptosis.

**Wider implications of the findings:** Measures of ovarian hormones should be rigorously included in future studies assessing cf-miRNA expression in females and used as time-varying confounders.

**Limitations, reasons for caution:** Our study was conducted on a relatively small cohort of patients. However, it was tightly controlled for endogenous and exogenous confounders, which is critical to ensure robust and reproducible cf-miRNA research.

**Wider implications of the findings:** Our results reinforce the importance of accounting for female-specific biological processes in physiology research by implementing practical or statistical mitigation strategies during data collection and analysis.

**Study funding/competing interest(s):** This study was supported by the clinique romande de réadaptation, Sion, Switzerland. Prof. Severine Lamon, is supported by an Australian Research Council Future Fellowship (FT10100278). The authors declare no competing interest

**Trial registration number:** N/A.

## Introduction

MicroRNAs (miRNAs) are short, non-coding RNA molecules encoded in the introns and exons (38) of the nuclear genome (14) and produced in most human tissues (24). MiRNAs are major regulators of gene and protein expression that directly or indirectly influence the translation process (12, 26, 46), primarily via the specific repression of target mRNA molecules (12).

Most physiological fluids including serum, plasma, urine, saliva and breast-milk contain a fraction of the miRNA pool (49). These extracellular miRNAs are referred to as cell-free or “circulating” miRNAs (cf-miRNAs). Cf-miRNAs are passively or selectively secreted by cells (3) into bodily fluids (49), and may contribute to cross-tissue communication by exerting their effects on recipient cells (11, 43). Although their biological role is not fully elucidated, many pathophysiological conditions, including a number of cancers (23), are associated with a dysregulation of the cf-miRNA profile in serum or plasma. Serum and plasma cf-miRNAs present the advantage of being relatively easy to access through venepuncture, making them potentially valuable biological markers (15).

Research into the circulating forms of miRNAs has rapidly evolved over the last decade and methodological approaches have become more stringent (15). In an attempt to reduce participant variability, most studies have focused on male only-cohorts (10, 33). There is however a global call to include sex as a biological variable in fundamental and applied research studies to broaden their range of applicability (5, 10, 33). Increasing evidence suggests that the regulation of miRNA expression is sex-specific (40, 41) and that males and females display different miRNA profiles in a variety of tissues and disease states (13, 40), including in plasma (30). Our group recently suggested that, in parallel to male-female comparisons, researchers should also investigate the effect of female sex hormone fluctuations due to the menstrual cycle and/or oral contraception on their physiological outcome of interest (19, 22). This holds true for molecular outcomes, including miRNAs, whose expression levels in reproductive tissues fluctuate throughout the menstrual cycle (47).

Evidence around the effect of female sex hormones on the plasma cf-miRNA profile is limited. In females with polycystic ovary syndrome (PCOS), presenting with higher levels of androgen hormones and lower levels of oestradiol when compared to healthy controls, the expression levels of 15 cf-miRNAs were closer to the average male than to the average healthy female expression levels (30). This dysregulation suggests a pattern of androgenisation associated to sex steroid hormones (30) and warrants the investigation of the effect of natural female hormone fluctuations on the cf-miRNA profile. An early human study investigating cf-miRNA expression at four time points across the menstrual cycle (37) suggested that the overall cf-miRNA profile remains stable throughout the cycle. However, no individual data or analysis were made available. In addition, this study failed to control for differences in RNA extraction and reverse-transcription, which considerably increase between-sample variability (15). In contrast, a study conducted in larger mammals (Holstein-Friesian heifers) identified a series of cf-miRNAs whose expression levels varied during the bovine oestrous cycle (17).

There is an increasing body of research surrounding the role and regulation of cf-miRNAs (15) and their potential to be used as non-invasive, cheap biological markers for a number of disorders including cancers (23), cardiovascular (52) and liver disease (42). This should prompt researchers to broaden the field of applicability of their findings by understanding the effect of the female hormonal environment. This study aims at taking the first step by conducting a stringent, tightly controlled, high-throughput pilot analysis of the plasma cf-miRNA profile across the female menstrual cycle.

## Methods

### Study population

Twenty eumenorrheic biological females (defined as a person born with XX chromosomes) were recruited among health-care providers of a rehabilitation clinic in Switzerland. Females aged 18-50 and having had a regular menstrual cycle during the last 18 months were invited to take part in the study. Exclusion criteria included hormonal contraception (including oral contraceptives, implant and hormonal IUD); pregnancy or attempting to get pregnant; breastfeeding; gynaecological conditions; history of hepatitis B, C or D; HIV infection. General health characteristics, menstrual anamnesis, use of medication and health events occurring during the menstrual cycle (e.g. menstrual pain, abnormal uterine bleeding or headache) were recorded. Four participants were excluded from the study: two females started hormonal contraception during the sample collection period, and the hormones levels of two other females could not be matched with a specific phase of the menstrual cycle (9). Sixteen women were therefore included in the final analysis. The study protocol was approved by the local ethics committee (CER-VD 2018-01914) and conducted according to the recommendations of the *Declaration of Helsinki* (1) and its later amendments. Informed, written consent was collected from each participant.

### Procedures

All participants had been monitoring their menstrual cycle using their mobile phone app of choice for at least six months before entering the study to ascertain the regularity of their menstrual cycle. Blood sampling was scheduled at three time-points corresponding to the early follicular phase (T1, ideally defined as the second day of the menstruation), the ovulation phase (T2, ideally defined as one day past the luteinizing hormone (LH) surge) and the mid-luteal phase (T3, ideally defined as seven days past T2). Figure 1 depicts the outline of the study. Immuno-chromatographic, urine-based LH detection tests (*Livsane LH20M, Pharmapost AG, Switzerland*) were used to identify the LH peak according to the manufacturer’s instructions. Menstrual cycle phase identification was conducted according to the recommendations from Elliott-Sale et al. (9) modified to account for the ovulation test result and the surge in LH and FSH (Supplementary Table I). When a resolution could not be met, data were compared to three different clinical reference ranges for oestradiol and progesterone according to (9). Data that did not comply with this framework were not considered, resulting in the exclusion of two individual data points.

**Figure 1.**
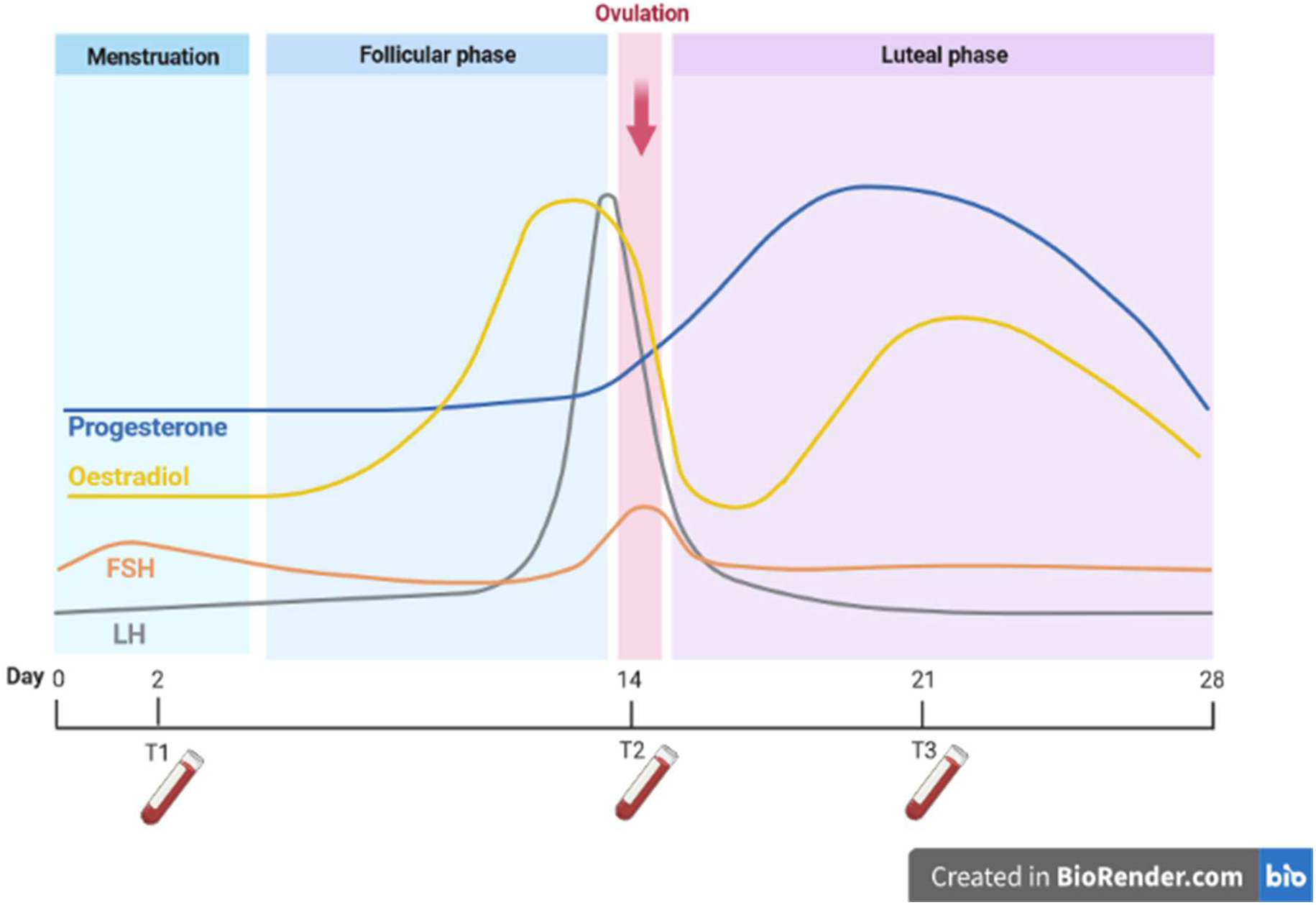
Outline of the study. Blood samples were collected at three time points of the menstrual cycle. T1 = early follicular phase, ideally defined as the second day of the menstruation; T2 = ovulation phase, ideally defined as one day past the luteinizing hormone (LH) surge; T3 = mid-luteal phase T3, ideally defined as seven days past T2. Created with BioRender.com.

### Blood collection

Peripheral blood samples were collected from the antecubital vein after an overnight fast in 2 × 7.5 mL EDTA and 1 × 7.5 mL Serum-Gel tubes (*S-Monovette®, Sarstedt International, Numbrecht, Germany*) using a butterfly device. The tubes were immediately centrifuged at 3’000 g at 4°C for 10 min. Plasma and serum were aliquoted in 1.5 ml RNase-free and DNase-free tubes and stored at −80°C until further processing. Samples with visible signs of haemolysis were excluded from the analysis.

### Hormone analysis

Serum levels of ovarian hormones estrogen, progesterone, luteinizing hormone (LH) and follicle-stimulating hormone (FSH) were measured by electrochemiluminescence using an Elecsys & Cobas system (*Roche, Rotkreuz, Switzerland*) at the Clinical Chemistry and Toxicology Department of the Central Institute of the Hospital of Valais (ICHV), Sion, Switzerland.

### RNA isolation and cDNA synthesis

Prior to RNA isolation, plasma samples were centrifuged for 5 min at 3’000 g. Then, 300 μl of supernatant was transferred into a new tube for RNA extraction using the miRNeasy^®^ Serum/plasma advanced kit (*Qiagen, Venlo, The Netherlands*) following the manufacturer’s instructions. To control for the reproducibility of the extraction, 3, 0.03 and 0.00003 fmol of UniSp2, UniSp4 and UniSp5 (*Qiagen*), respectively, were added to the lysis buffer of each sample. The total RNA fraction was eluted in 20 μl RNase-free water. Two uL of total RNA was used to synthetized cDNA using the miRCURY LNA RT kit (*Qiagen*) according to the manufacturer’s protocol. 0.002 fmol of the synthetic miRNA cel-miR-39-3p (*Qiagen*) and 0.15 fmol of the spike-in UniSp6 (*Qiagen*) were added to the cDNA to control for the reproducibility of the cDNA synthesis.

### miRNA profiling

Plasma miRNA expression analysis was performed using the Human serum/plasma Focus, miRCURY LNA miRNA Focus PCR panel (*Qiagen*). This sensitive and specific PCR-based method measures the expression levels of 179 serum-enriched miRNAs and includes internal and external quality controls. These miRNAs have been carefully selected based on their expression in plasma samples reported by previous studies and manufacturer recommendations. Amplification was performed with the LightCycler 480 Real-Time PCR system (*Roche*) as previously described (8).

### Data quality assessment

After the initial visual inspection, haemolysis was assessed using the difference in cycle quantification value (Cq) between hsa-miR-23a-3p, which is unaffected by haemolysis, and hsa-miR-451a, which is highly expressed in red blood cells (7). Based on results from the literature, a difference smaller than seven Cq was considered acceptable (7). According to this criteria, two samples displayed faint signs of haemolysis and were excluded from the analysis.

As recommended by the Biofluids Guidelines from Qiagen and described in (31), a pool of exogenous spike-in controls UniSp2, UniSp4 and UniSp5, and a pool of exogenous spike-in controls UniSp6 and cel-miR-39-3p were added prior to the RNA isolation and cDNA synthesis steps, respectively. Six replicates of UniSp3, an exogenous quality control for PCR amplification, were included in the pre-coated PCR panel and UniSp3 was used as an inter-plate calibrator (see data processing below). The between-sample coefficients of variations (CVs) for UniSp2 (4.44 %), UniSp4 (3.45%), UniSp5 (3.02%), and UniSp6 (2.06%) were considered excellent.

### Data processing and normalization

Cq values were calculated by applying the second derivative method and transformed into arbitrary units using the formula (2^Ct) × (10^10). Melting curves were examined automatically using the LightCycler 480 software version 1.5.0, and individual data points were manually excluded if a double peak or a “shoulder” was visible, leading to the exclusion of 181 data points (1.97%). Six miRNA datasets presented an average Cq value > 35 and were excluded from further analysis. Following this, all individual Cq values > 35 remaining in the dataset were excluded from further analysis, leading to the exclusion of an additional 189 data points (2.29%). Of the remaining miRNA datasets, eleven (6.45%) missed more than 20% of their individual data points and were excluded on this basis. For the remaining 157 miRNAs, the arithmetic mean of the transformed Cq values (arbitrary units) of the six endogenous UniSp3 quality controls was calculated for each plate and allocated a random value of 1 to account for inter-plate variability. Finally, the geometric mean of the transformed Cq values (arbitrary units) of all miRNAs considered for the final stage of the analysis was used for global normalization.

### Statistical analysis

All data were analysed using R studio 4.1.3 (35). Missing data were imputed using the *mice* package (45) and iteration three was randomly selected for all analyses. To meet the statistical assumptions of the regression (normally distributed residuals), ovarian hormone levels (oestrogen, progesterone, LH and FSH) were log-transformed. To identify changes in cf-miRNA expression across the menstrual cycle, we used linear mixed models for repeated measures as implemented in the *variancePartition* package in R (16). Participant ID was used as the random effect to account for repeated measures and all models were adjusted for age. The model was of the form: *miRNA* ~ *Timepoint* + *Age* + (1|*ID*). The model was then fitted with contrasts to examine changes in miRNA expression between the different phases of the menstrual cycle. We then added ovarian hormone (oestrogen, progesterone, LH and FSH) concentrations into the model to understand if hormones explained any of the variability in cf-miRNA expression. Due to co-linearity, models of the form: *miRNA* ~ *Hormone* + *Timepoint* + *Age* + (1/*ID*) were run separately for each hormone. We used the Benjamini-Hochberg adjustment to correct for multiple comparisons to minimize the risk of false positive results. However, given the exploratory nature of the study, we decided to report both unadjusted and adjusted *p values* for all analyses.

Time series clustering analysis was performed to identify similarities in expression patterns across the menstrual cycle. The elbow method and gap statistics were used to identify the optimal number of clusters (N = 3) for analysis. The cf-miRNAs were grouped based on similarities in their slopes regardless of the distance (level of expression) between them using the *tSCR* R package (28). Gene target analysis was conducted using the R package *multi-MiR* (39). Only gene targets identified via luciferase reporter assay or western blotting were included in the gene target and pathway enrichment analysis. Over representation analysis (ORA) was used for gene target enrichment and implemented by the R package *clusterProfiler* (53). The background gene list used for the ORA analysis comprised all validated targets of the c-miRNAs detected in the PCR panel, using the same stringency criteria as above. Tissue enrichment of the validated gene targets was conducted in the R package *TissueEnrich*. Finally, transcription factor-microRNA interactions were explored using TransmiR v2.0 (44) and compared using two-sample tests of proportion ran in Stata 16.0. *P values* were deemed statistically significant when *p* <0.05. Data are presented as Mean ± SD unless stated otherwise. The following packages were also used in our analysis; lme4 (4), lmerTest (21), and tidyverse (51). The full R code can be found at https://github.com/DaniHiam/menstrual-cycle-circulating-microRNAs.

## Results

### Participant characteristics

Sixteen, eumenorrheic biological females aged 37.8 ± 6.8 years participated in the study. Participants were apparently healthy with a BMI of 22.3 ± 3.4. None of the participants smoked regularly and two participants smoked occasionally. All participants had a regular menstrual cycle for at least six months prior to entering the study. Table 1 recapitulates participant hormone levels at each sample collection point.

**Table 1.**
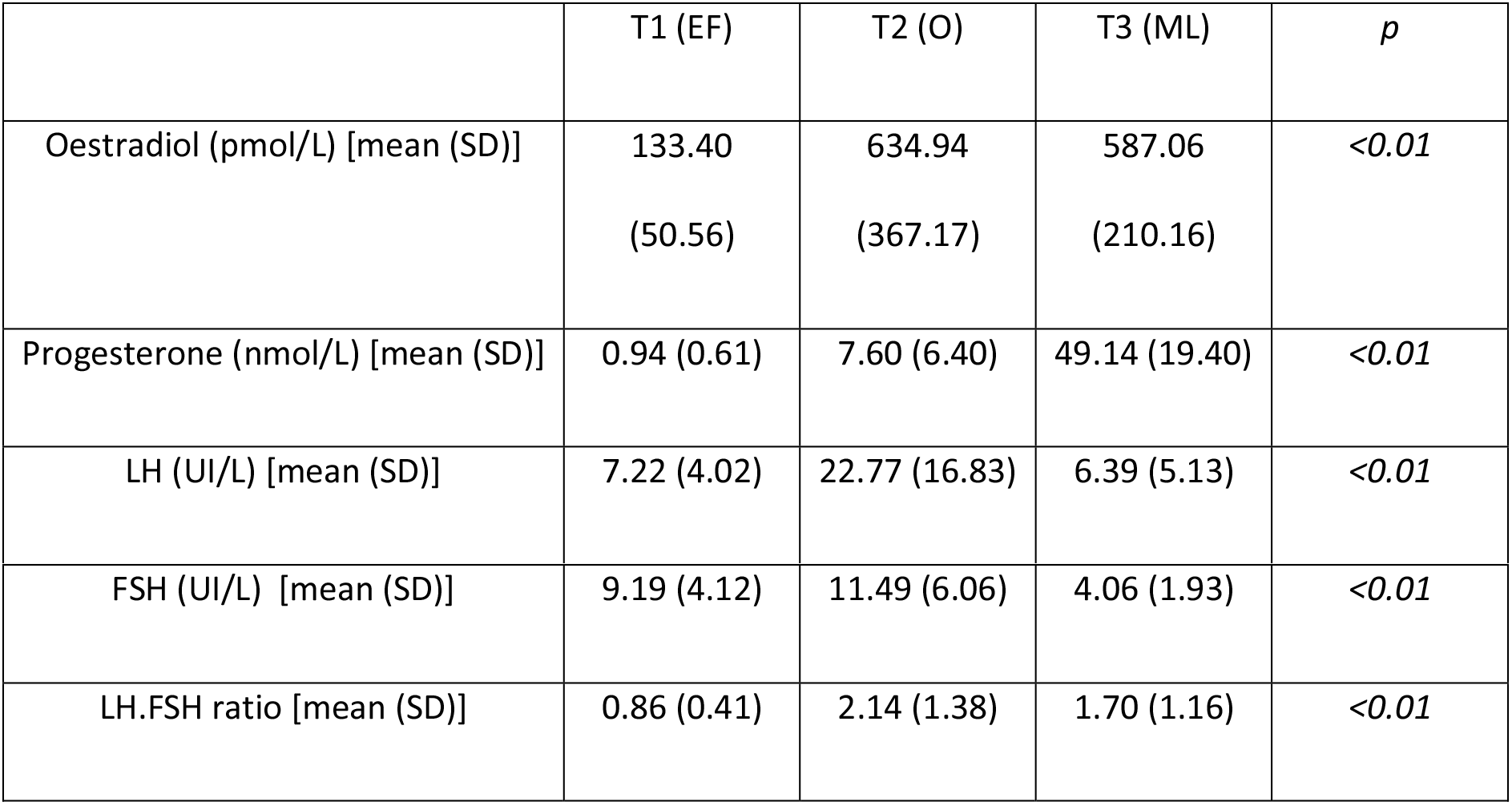
Hormone levels at each blood collection point. T1 = early follicular (EF) phase, ideally defined as the second day of the menstruation; T2 = ovulation (O) phase, ideally defined as one day past the luteinizing hormone (LH) surge; T3 = mid-luteal phase T3 (ML), ideally defined as seven days past T2.

|  | T1 (EF) | T2 (O) | T3 (ML) | *p* |
| --- | --- | --- | --- | --- |
| Oestradiol (pmol/L) [mean (SD)] | 133.40<br>(50.56) | 634.94<br>(367.17) | 587.06<br>(210.16) | $<0.01$ |
| Progesterone (nmol/L) [mean (SD)] | 0.94 (0.61) | 7.60 (6.40) | 49.14 (19.40) | $<0.01$ |
| LH (UI/L) [mean (SD)] | 7.22 (4.02) | 22.77 (16.83) | 6.39 (5.13) | $<0.01$ |
| FSH (UI/L) [mean (SD)] | 9.19 (4.12) | 11.49 (6.06) | 4.06 (1.93) | <0.01 |
| LH.FSH ratio [mean (SD)] | 0.86 (0.41) | 2.14 (1.38) | 1.70 (1.16) | <0.01 |

### The effect of the menstrual cycle on circulating miRNA expression

Of the 174 endogenous, plasma-enriched miRNAs included on each plate, 157 (92%) met our minimal expression criteria and were considered for further analysis. An unsupervised, clustered heat map of the z-scored expression of all analysed miRNAs grouped by time point is presented in Figure 2.

**Figure 2.**
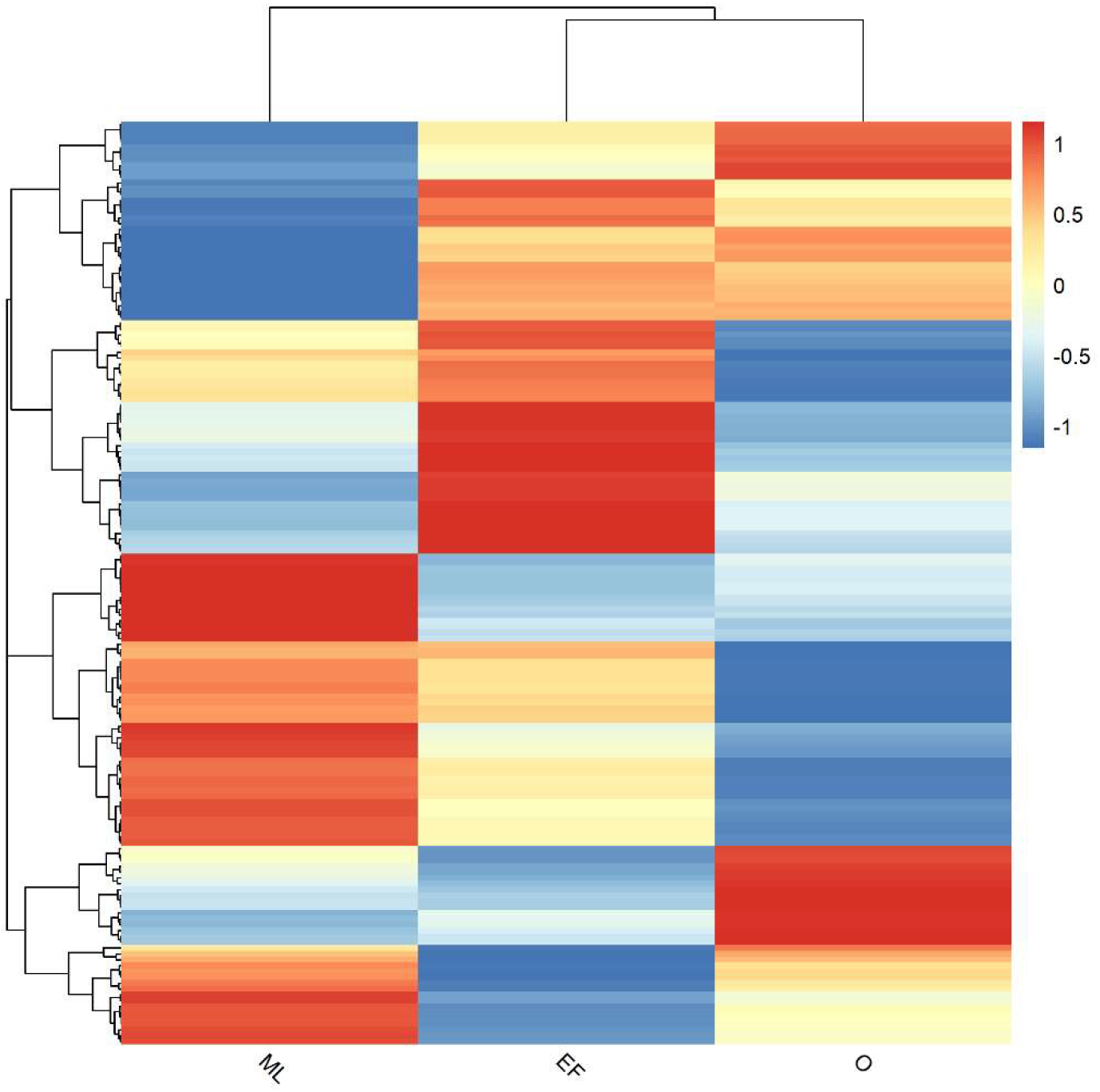
Unsupervised clustered heat map of the z-scored expression of 157 plasma-enriched miRNAs meeting the minimal expression criteria in the mid-luteal (ML), early follicular (EF), ovulatory (O) phases of the menstrual cycle (N=16).

The expression levels of six individual miRNAs, hsa-miR-99b-5p, hsa-miR-1260a, hsa-let-7e-5p, hsa-miR-30c-5p, hsa-miR-766-3p and hsa-miR-326, displayed significant differences with the menstrual cycle (*p<0.05*) (Figure 3). Contrasts were then added in the model. Additional miRNAs presented significant changes between individual time points (T1 vs T2: hsa-miR-30e-5p, hsa-miR-155-5p, hsa-miR-27a-3p, hsa-miR-382-5p, hsa-miR-590-5p, hsa-miR-374a-5p, hsa-miR-32-5p, hsa-let-7d-5p; T1 vs T3: hsa-miR-30c-5p; T2 vs T3: hsa-miR-374a-5p, hsa-miR-454-3p, hsa-miR-32-5p, hsa-miR-590-5p, hsa-miR-30e-5p, hsa-miR-424-5p, hsa-miR-126-5p, hsa-miR-1260a, hsa-miR-154-5p, hsa-miR-136-3p, hsa-miR-324-5p, hsa-miR-382-5p, all *p<0.05*). Since our study is exploratory, we elected to report non-adjusted *p values. P values* adjusted for multiple comparisons are available in Supplementary Data 1. Between-participant variability explained a greater portion of the variance than any other variable (Supplementary Figure 1), indicating that, in future studies, a larger cohort may help overcoming the issue of small effect sizes.

**Figure 3.**
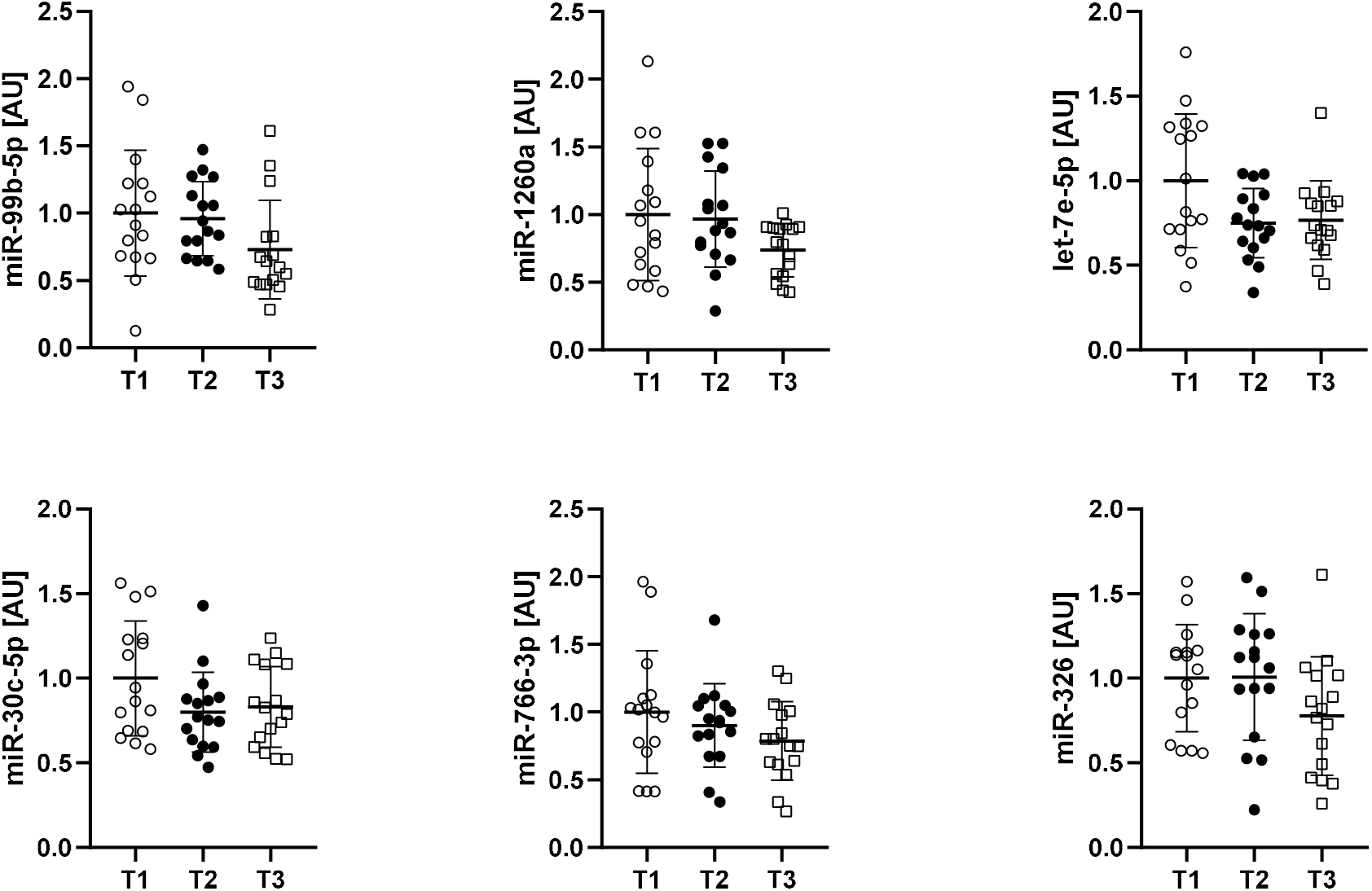
Individual miRNA levels of hsa-miR-99b-5p, hsa-miR-1260a, hsa-let-7e-5p, hsa-miR-30c-5p, hsa-miR-766-3p and hsa-miR-326 significantly varied across the menstrual cycle (all *p<0.05*). T1 = early follicular (EF) phase, ideally defined as the second day of the menstruation; T2 = ovulation (O) phase, ideally defined as one day past the luteinizing hormone (LH) surge; T3 = mid-luteal (ML) phase T3, ideally defined as seven days past T2. Values were normalized using the global normalization method and displayed as Mean ± SD (N=16).

In an attempt to identify similarities in expression patterns across the menstrual cycle, three clusters containing 26, 4 and 127 miRNAs, respectively, were identified based on similarities in their slopes (28). None of these clusters however displayed significant changes in miRNA expression patterns across time (Supplementary Figure 2). Instead, we ran gene target prediction and pathway analysis on the six above-mentioned miRNAs that significantly varied with time in our exploratory analysis to identify potential common biological pathways. Tissue enrichment was conducted on the premises that each miRNA gene target had been experimentally confirmed either by luciferase reporter assay or Western Blot. Gene targets were significantly enriched in the endometrium, the cervix and uterine tissue, the placenta and the ovary (all *p<0.05*) (Figure 4). This analysis was then repeated with the complete list of miRNAs presenting significant contrasts between any two time points and yielded similar results (Supplementary Figure 3).

**Figure 4.**
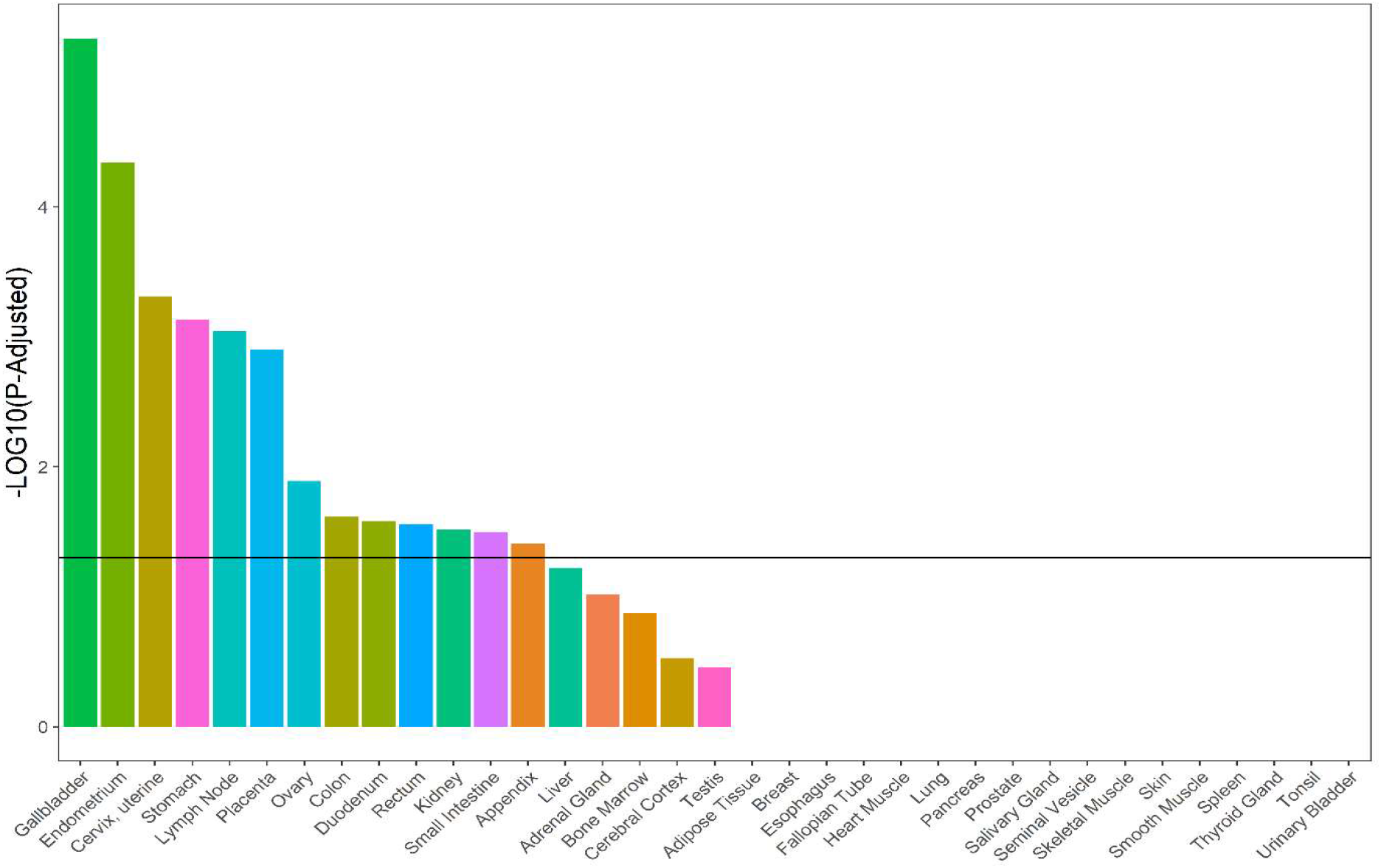
Tissue enrichment of the validated gene targets of the cf-miRNAs varying across the menstrual cycle. Solid black line indicates significance (*p < 0.05*).

Hierarchical clustering of enriched terms was conducted on the same list of genes. The most significantly enriched biological pathways pertained to cellular responses to development. Amongst the four pathways demonstrating the highest number of gene targets and levels of statistical significance were epithelium development; regulation of organismal development; positive regulation of cell population proliferation; and positive regulation of cell death.

**Figure 5.**
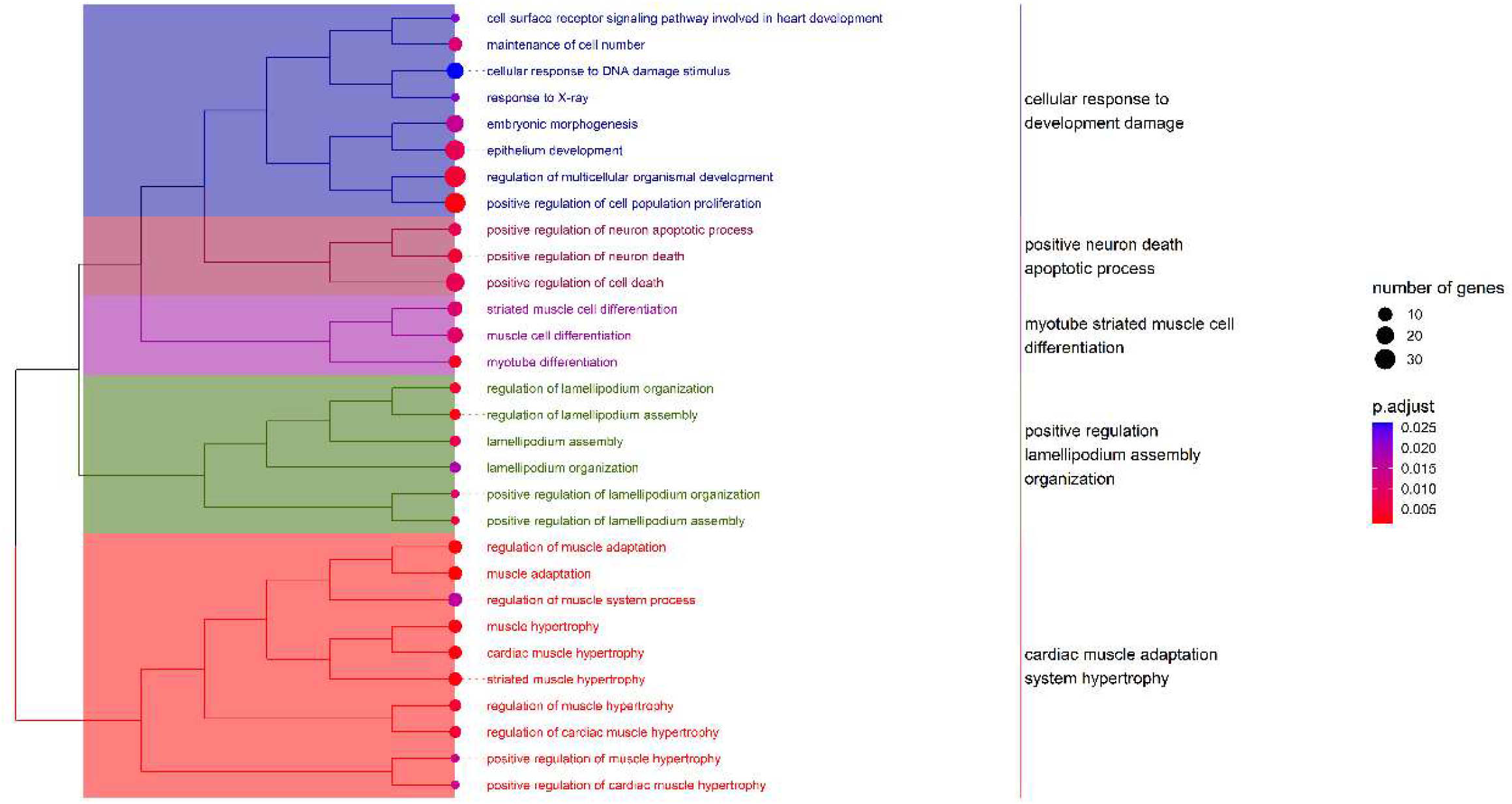
Hierarchical clustering of enriched terms for the validated gene targets of the cf-miRNAs varying across the menstrual cycle.

### The effect of female sex hormones on circulating miRNA expression

We then investigated whether plasma levels of oestradiol, progesterone, LH and FSH explained any of the variance in miRNA expression across time once adjusted for age. The expression levels of 48 individual miRNAs were significantly associated with progesterone, oestradiol, LH and/or FSH levels prior to FDR adjustment (all *p<0.05*) (Figure 6). *P values* adjusted for multiple comparisons are available in Supplementary Data 2.

**Figure 6.**
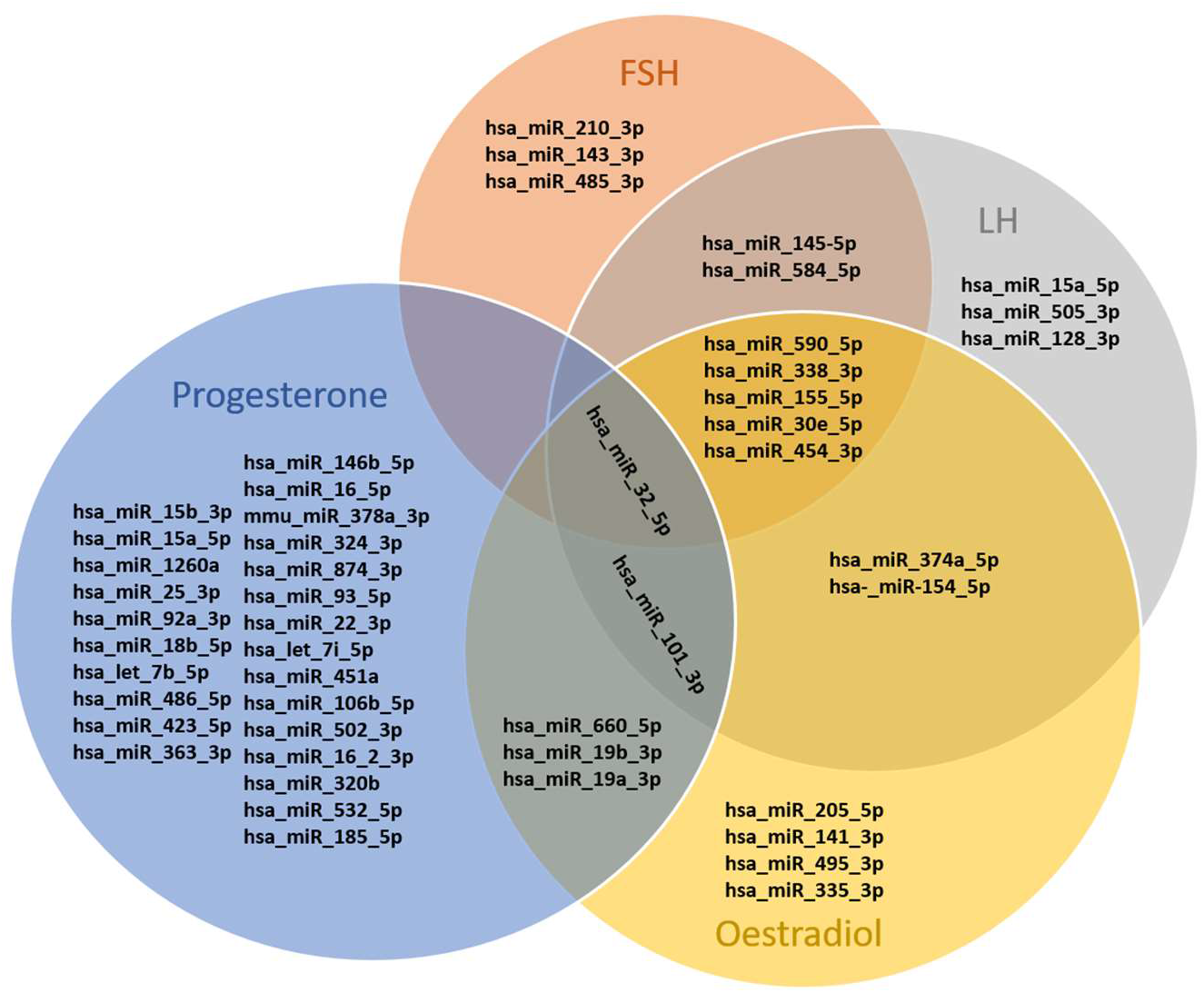
Individual miRNAs that were significantly associated with progesterone (30 miRNAs, blue), oestradiol (16 miRNAs, yellow), LH (14 miRNAs, grey) and/or FSH (11 miRNAs, orange) plasma level in age-adjusted regression models.

Finally, we investigated whether specific transcription factors were more likely to regulate the expression of the 48 miRNAs associated with progesterone, oestradiol, LH and/or FSH levels. Numerous transcriptions factors presented a significant interaction with the 48 miRNAs of interest (*p<0.05*), but none of them was specific to any of the female sex hormones. Only three transcription factors, the Aryl hydrocarbon receptor nuclear translocator (ARNT), the Myb proto-oncogene protein (MYB) and Upstream stimulatory factor 1 (USF1) were significantly over-represented as regulators of the list of miRNAs of interest when compared to the complete list of miRNAs detected in the PCR panel (*p<0.05*) (Supplementary Data 2).

## Discussion

Here we report the results of an exploratory study investigating the effects of the menstrual cycle on plasma cf-miRNA expression in 16 eumenorrheic females. One prior study conducted in nine young females suggested that the overall cf-miRNA profile remained stable throughout the menstrual cycle. However, this study critically failed to control for differences in RNA extraction and reverse-transcription, and individual data or analysis were not made available (37). A more recent study investigating a suite of serum cf-miRNAs as potential markers for endometriosis found that the phase of the menstrual cycle, which was recorded but not controlled for, did not influence the expression levels of the six miRNAs used in their diagnosis algorithm (29). Again, differences in RNA extraction and reverse-transcription were not accounted for. In contrast, our study, while conducted on a small cohort, was tightly controlled for endogenous and exogenous confounders, which is critical to ensure robust and reproducible cf-miRNA research (15). Our data highlight numerous associations between phases of the menstrual cycle, female sex hormone levels and plasma cf-levels.

The idea that sex hormones influence miRNA expression in human tissue is not new (32), but little research has focused on the menstrual cycle and its influence on cf-miRNA levels, maybe because of the intrinsic complexity of these two systems (15, 22). In larger mammals, Ionnadis et al (10) reported a series of plasma cf-miRNAs whose expression levels varied during the estrous cycle of Holstein-Friesian heifers. Interestingly, we identified two conserved miRNAs, hsa-miR-99b-5p (homologous to bta-miR-99b) and hsa-miR-155-5p (homologous to bta-miR-155) that also displayed significant contrasts between time points in humans. While these contrasts did not retain their significance once adjusted for FDR in either study, the pattern of change remained similar between both species. The expression levels of hsa-miR-99b-5p decreased by 27 % (humans) and 25% (bovines) (10), respectively, in the mid-luteal phase of the menstrual cycle when compared to the early-follicular phase. MiR-99b-5p is a well-studied miRNA that is mechanistically associated with the AKT/MTOR pathway and a direct target of MTOR, RPTOR and IGF1 3’-UTRs (54). This pathway retains its homology in the mouse and mediates oestradiol-induced protein synthesis in mouse reproductive tissue (48). In humans, miR-99b clusters with hsa-let-7e on chromosome 19, which also demonstrated a 25% decrease between the early-follicular and mid-luteal phases. Supporting a role for this cluster in human reproductive tissues, hsa-let-7e expression significantly increased by about 2-fold when treated with oestradiol, but not progesterone, in human endometrial stromal cells (36).

Similarly, the expression levels of hsa-miR-155-5p increased by 45% (humans) and 32% (bovines) (10) in the ovulatory when compared to the early-follicular phase. MiR-155-5p is a well-studied oncomiR that is upregulated in different types of breast cancers (6, 18, 20) and associated with reduced expression of the estrogen receptor and progesterone receptor in human primary breast tumors (34).

Further relevant comparisons can be drawn with the recent pilot study by McGregor et al. describing the fluctuations of miRNAs in human adipose tissue at four points of the menstrual cycle (25). The time points used in the McGregor study do not fully align with ours (they collected a late-follicular and a post-ovulatory time point, while we collected a peri-ovulatory time point instead), but some of the miRNAs differentially expressed in adipose tissue were also differentially expressed in plasma, again with the limitation of FDR adjustment. Surprisingly, we report opposite patterns for hsa-miR-155-5p, which tends to increase around ovulation in plasma, but to decrease in adipose tissue, and hsa-miR-30c-5p, which tends to increase during the mid-luteal phase in plasma but to increase in adipose tissue. Cf-miRNAs are passively or selectively secreted by cells (3) into bodily fluids (49). It is therefore possible that adipose tissue acts as a reservoir for a pool of miRNAs that are secreted into plasma at specific time points to exert their effect on recipient cells (11, 43).

The gene targets of the miRNAs fluctuating with the menstrual cycle were significantly enriched within the female reproductive tissues endometrium, cervix, uterine, placenta and ovary. Despite an exponential increase in circulating miRNA expression research over the last decade (15), insights into the role and regulation of these molecules remain elusive. How specific cf-miRNA transport processes are is unclear, and the mechanisms governing their uptake by recipient cells remain mostly unknown. A number of studies suggest highly selective and tightly regulated processes (2, 43, 50), which may underpin a whole level of cross-tissue communication (11, 50). Our findings support this hypothesis, where the menstrual cycle may control the tissue-specific secretion of cf-miRNAs targeting the female reproductive apparatus. In addition, the gene targets of these cf-miRNAs primarily regulated the regulation of epithelium development. Endometrial epithelial cell proliferation peaks in the follicular or “proliferative” phase of the menstrual cycle, in preparation for a potential implantation, while apoptosis occurs later in the luteal or “secretory” phase (27). These results indicate that cf-miRNA may be intrinsic regulators of these processes in female reproductive tissues. This regulation may however not specifically result from sex-hormone specific transcription factors, such as the oestrogen receptors (ESR1 and ESR2) or the progesterone receptor (PR), acting on miRNA transcription. Future studies may therefore focus on investigating indirect mechanisms.

Our study is exploratory. The one prior study reporting human cf-miRNA expression across the menstrual cycle did not make individual data or analysis available (37). To the best of our knowledge, no other suitable data set was available in the literature to base our power calculation on. We therefore established our sample size based on feasibility as well as previous, similar human studies conducted in different tissues (25). Bearing the restrictions of this approach in mind, no participant was added to the original cohort past the end of data collection. Our study was further limited by the stringent Benjamini-Hochberg adjustment, which was applied to our *p-values* to account for the multiple cf-miRNAs measured on the same assay plate. While none of the adjusted *p-values* reached statistical significance, unadjusted *p-values* indicate that our study would have been adequately powered for more targeted analysis and may therefore be considered as hypothesis-generating. Unadjusted *p-values* suggest a range of likely associations between cf-miRNA expression and serum sex hormone levels. We therefore recommend that future studies investigating the individual or collective expression levels of miRNAs in female plasma should 1) include measures of serum progesterone, oestrogen, FSH and LH levels, and 2) adjust their mixed-models by accounting for these hormone levels as time-varying confounders. The full code based on R language is available for use at https://github.com/DaniHiam/menstrual-cycle-circulating-microRNAs.

In conclusion, we report the results from a pilot study investigating the effects of the menstrual cycle on the plasma cf-miRNA expression profile in humans. We found numerous associations between cf-miRNAs levels, phases of the menstrual cycle and ovarian hormone levels prior to adjusting our results for multiple comparisons. Stringent bioinformatics analysis indicates that the menstrual cycle may regulate the expression of cf-miRNAs targeting cell development and proliferation in female reproductive tissues. Our results reinforce the importance of accounting for female-specific biological processes in physiology research by implementing practical or statistical mitigation strategies during data collection and analysis.

## Authors’ roles

Severine Lamon: conception and design, analysis and interpretation of data, drafting the article, final approval of the version to be published

Joane Le Carré : conception and design, acquisition of data, critical revision for important intellectual content, final approval of the version to be published

Giuseppe Petito : acquisition of data, critical revision for important intellectual content, final approval of the version to be published

Hong Phuoc Duong : analysis and interpretation of data critical revision for important intellectual content, final approval of the version to be published

François Luthi : analysis and interpretation of data, critical revision for important intellectual content, final approval of the version to be published

Danielle Hiam : analysis and interpretation of data, drafting the article, final approval of the version to be published

Bertrand Léger : conception and design, analysis and interpretation of data, critical revision for important intellectual content, final approval of the version to be published

All authors agree to be accountable for all aspects of the work in ensuring that questions related to the accuracy or integrity of any part of the work are appropriately investigated and resolved

## Acknowledgments

Severine Lamon is supported by a Future Fellowship from the Australian Research Council (FT210100278). Danielle Hiam is supported by an Executive Dean’s Postdoctoral Research Fellowship from Deakin University.

